# Atorvastatin-induced vascular weakness models brain haemorrhage in vascularised cerebral organoids

**DOI:** 10.64898/2026.04.20.719465

**Authors:** Siobhan Crilly, Sugunapriyadharshini Sundararaman, Michael J. Haley, Elisa Segantin, Nicola Campbell, Eulalie J. Lafarge, Antonn Cheeseman, Josep Fumadó Navarro, Declan McKernan, Kevin N. Couper, Mihai Lomora

**Affiliations:** School of Biological and Chemical Sciences, College of Science and Engineering, University of Galway, Galway, Ireland; CÚRAM, Research Ireland Research Centre for Medical Devices, University of Galway, Galway, Ireland; Institute for Health Discovery and Innovation, University of Galway, Ireland; Lydia Becker Institute of Immunology and Inflammation, Faculty of Biology, Medicine & Health, Manchester Academic Health Science Centre, University of Manchester, Oxford, Manchester, M13 9PT, United Kingdom; Geoffrey Jefferson Brain Research Centre, Faculty of Biology, Medicine & Health, Manchester Academic Health Science Centre, University of Manchester, Oxford, Manchester, M13 9PT, United Kingdom; Bioimaging Core Facility, Faculty of Biology, Medicine and Health, University of Manchester, Manchester, M13 9PT, United Kingdom; Pharmacology and Therapeutics, School of Pharmacy and Medical Sciences, College of Medicine, Nursing and Health Sciences, University of Galway, Ireland

**Keywords:** Stroke, intracerebral haemorrhage, atorvastatin, brain organoids, blood-brain barrier, cholesterol

## Abstract

Intracerebral haemorrhage is the most severe subtype of stroke; however, pre-clinical investigation often fails to translate to the clinic. Cerebral organoids offer an adaptable, in vitro model of human brain tissue for pre-clinical investigation of disease. We recently demonstrated that the tissue can be successfully vascularised to mimic the cerebrovasculature.

Cerebrovascular weakness was induced with atorvastatin to mimic damage observed in intracerebral haemorrhage and to replicate the disease’s pathological features. We used atorvastatin to disrupt functional morphology in human brain microvascular endothelial cells in 2D and 3D model systems. Whole human blood was added to initiate damage to cerebral tissues.

Vascularised cerebral organoids exhibited loss of vascular integrity when treated with atorvastatin. Tissue was vulnerable to injury from human whole blood, and an innate immune response was initiated, resulting in increased cell death.

Here we show that vascularised cerebral organoids demonstrate a novel model platform for investigating pathology associated with human whole blood insult in intracerebral haemorrhage.

## Introduction

Intracerebral haemorrhage (ICH) or haemorrhagic stroke is a leading cause of death and disability worldwide.^1^ Determining pathological mechanisms and therapeutic targets depends on preclinical modelling. Recently, the array of models available to recreate aspects of clinical disease has expanded.^2,3^ However, options for interrogating living human tissues remain limited. Haemorrhagic stroke is a multi-systemic disorder caused by a rupture in the cerebrovasculature and blood leaking into the brain parenchyma, and stimulating both a localised and peripheral innate immune response.^4^ Advances in 3D cultures in biomaterials and organ-on-chip applications have enabled the development of a blood-brain barrier (BBB) model that can inform understanding of mechanisms of vascular pathology. Although these models have been useful to determine developmental disease processes,^5^ there has been no application of such complex models to ICH research.

Recently, culturing 3D organoids from human stem cells has accelerated the capacity of multi-cell cultures in vitro to recreate complex tissues. Cerebral organoids have been derived to recreate brain tissue and are often heterogenous cultures of different tissue types. Directional differentiation protocols have been developed to create specific brain regions.^6^ They have been explored as a tool for modelling ischaemic stroke pathology^7^ and are relatively easy models for exploring oxygen-glucose deprivation damage.^8^ Recently, we developed a strategy to incorporate human brain microvascular endothelial cells (HBMECs) to permeate the cerebral tissue and form vascular-like networks.^9^ Vascularised cerebral organoids exhibit CD31-positive endothelial cell networks within the tissue, neuroepithelial markers such as zonula occludens-1 (ZO-1) and functional tight junction proteins such as VE-Cadherin and claudin5, the presence of pericyte and astrocyte cell markers, and less apoptosis within the organoid tissue compared to non-vascularised controls.

To recreate the vessel damage observed in ICH pathophysiology, we have investigated the efficacy of atorvastatin at disrupting endothelial cell function in both 2D in vitro models and 3D vascularised cerebral organoids. Previously, atorvastatin has been used to disrupt neurovascular development in *Danio rerio*^10^ and this resulted in stroke-like pathology in developing zebrafish larvae.^11,12^ Atorvastatin inhibits mevalonate, the rate-limiting enzyme in cholesterol biosynthesis, an essential process in neovascular development and crucial for tight junction protein anchoring in the formation of new blood vessels. Atorvastatin has also been shown to disrupt tight junction expression by abnormal protein phosphorylation.^13^ We sought to determine if atorvastatin can recreate functional damage to HBMECs, similar to that previously observed in zebrafish, and weaken the vascular network within a vascularised cerebral organoid to ultimately recreate ICH pathology within the system. This is the first time a vascularised cerebral organoid platform has been used to model haemorrhagic stroke and can provide insights into human pathology and recovery mechanisms, with applications in therapeutic screening in the future.

## Materials and methods

### Ethical approval

Ethical approval was provided by the University of Galway Research Ethics Committee for studies performed using whole human blood samples (REC Application Reference Number: 2025.05.00).

### Cell culture

Primary human brain microvascular endothelial cells (HBMECs) (Innoprot P10361) were maintained on 1% gelatin coated flasks in endothelial cell growth medium MV (PromoCell C-22020). Cells were maintained at 37°C, 5% CO2, and passaged at 70-80% confluency. Cells for experiments were used between passages 7 and 14. When cells were incubated in rotating culture, 6-well plates were placed on an orbital shaker set to 70 rpm.

Human induced pluripotent stem cells (iPSCs) (Gibco human episomal iPSC line A18945, female) were commercially obtained and maintained on Geltrex-coated ultra-adherent cultureware (Sarstedt Cell+) in Essential 8 Flex media (Gibco A2858501). Geltrex was prepared at a 1:100 dilution with Knockout DMEM media (Thermo Fisher, 10829018) and polymerised at 37°C for 2 hours. Cells were revived from liquid nitrogen storage and incubated with Essential 8 Flex media supplemented with 1X RevitaCell (Thermo Fisher, A2644501) for 24 hours. Every 3-4 days, cells were passaged 1:3 using Gentle Cell Dissociation Reagent (StemCell Technologies 100-0485) into freshly coated 6-well plates. Cells were stored in liquid nitrogen at passages 3 and 6 and used to generate organoids at passage 9 onwards.

### Cerebral organoid culture

Cerebral organoids were generated from iPSCs using the StemDiff cerebral organoid culture kit (StemCell Technologies 08570) adapted from published protocols^14^ with adaptation for vascularisation as previously reported.^9^ In brief, 9000 stem cells were added to low-adherence round-bottomed 96-well plates pre-rinsed with anti-adherence rinsing solution (StemCell Technologies 07010) on day 0 in embryoid body (EB) formation media with 10 μM Rho-associated coiled-coil containing protein kinase (ROCK) inhibitor (StemCell Technologies 72304). On days 2 and 4, EBs were fed with formation media without ROCK inhibitor. On day 5, EBs were transferred to a prepared low-adherence 24-well plate in induction media and incubated for 2 days until organoids had smooth, optically translucent edges. On day 7, organoids were embedded in 15 μl of 80% Geltrex diluted in endothelial cell media to expand the neuroepithelium. For vascularised organoids (V-COs), 50,000 HBMECs were suspended in the Geltrex prior to organoid embedding. Following 30 minutes at 37°C to polymerise the Geltrex, the organoids were washed into low-adherence prepared 6-well plates in expansion media. Organoids were cultured in 1:7 EC:expansion media supplemented with 50 ng/ml vascular endothelial growth factor (human recombinant VEGF-165, StemCell Technologies 78073). From day 10, the expansion medium was replaced with maturation medium, and the plate was placed on an orbital shaker at 70 rpm. Organoids were cultured until 40 days before experimental use.

### Drug treatment

Atorvastatin calcium trihydrate (Thermo Scientific 464810010) was reconstituted in DMSO (Merck, D2650-100ML) to generate a stock solution of 100mM. Stock solutions were diluted into warm media to generate the final working concentrations, and the maximum volume of DMSO was used as a vehicle control (1%).

### Tube formation assay

A 24-well plate was coated with 50 μL Geltrex, and then the matrix was polymerised at 37°C for 30 minutes. 40,000 cells were seeded in 0.5 mL, and the tubes were left to form for 24 hours. Atorvastatin was added either 1 hour after seeding to prevent tube formation or 24 hours after seeding to disrupt formed tubes. Images were taken using the EVOS M5000 digital inverted microscope, and network analysis was performed using AngioTool 2.0.^15^

### Cell viability assay

Supernatant was collected from adherent cell culture to test for cell viability using the CytoTox non-radioactive cytotoxicity assay (Promega G1780) as per the manufacturer’s instructions. In short, 50 μL of media was transferred to a flat-bottom 96 well plate and equal volume of the CytoTox regent was added. Cells were lysed using the kit lysis solution to generate a positive control. The plate was incubated at RT for 15 mins or until the positive control was dark red. Stop solution was added to each well and the absorbance at 492nm was recorded using BioTek multiwell plate reader (BioTek Epoch2) and Gen5 software (BioTek).

### Human blood

Heparinised human whole blood was obtained from the Irish Blood Transfusion Service and designated for research purposes. Blood from one donor was aliquoted and stored at -80°C until use.

### Filipin stain

Filipin (Medchem Express HY-N6716) was resuspended in DMSO to make a 5 mg/ml stock and diluted to 50 μg/ml into PBS for a working solution. After cells were fixed, filipin was added at the working concentration for 30 minutes in the dark. Cells were washed and imaged using a 405 nm emission. The Filipin-stained area was calculated using intensity analysis in ImageJ across 6 ROIs per well.

### ELISA assay

Supernatants from three batches of 40-day organoids after 4-hour incubation with human whole blood were diluted as required. Levels of IL-1β, IL-6, IL-8, IL-10, and TNF-α were quantified using standard sandwich ELISA protocols with human DuoSet kits (R&D Systems: DY206, DY208, DY210, DY201, DY217B), following the manufacturer’s instructions. Absorbance was measured at 450 nm, with correction at 540 nm, using a microplate reader (ELx800, BioTek). Cytokine concentrations were calculated by interpolating sample absorbance values from a standard curve using a cubic spline fit.

### RNA extraction, cDNA generation and qPCR analysis

Trypsinised cells or organoids were washed into 1.5ml Eppendorfs and lysed with TRIzol reagent (Thermo Fisher 15596026). Organoids were manually homogenised using a microfuge tube pestle. RNA was isolated using 75 μl of chloroform and centrifugation to isolate the aqueous layer. RNA was precipitated with isopropanol, pelleted, washed and resuspended in 20 μl RNAse-free water. RNA was purified with a DNase I (Invitrogen AM1906) incubation and quantified by NanoDrop 2000 Spectrophotometer. To generate complementary DNA sequence from isolated RNA, 800 ng of RNA was incubated with 1 µl of reverse transcriptase from the High-Capacity RNA-to-cDNA kit (Applied Biosystems 4387406) for 37 °C for 1 hour before reaction was terminated at 95 °C for 5 minutes. Gene expression analysis was performed using RT-qPCR using sample cDNA and SYBR Green Master mix (Applied Biosystems, 4367659) and primer sequences from Table 1. Relative quantities were normalised to the expression of housekeeper genes. Data was acquired using the Applied Biosystems QuantStudio3 and relative quantification evaluated from 2ΔΔC T ^16^.

**Table 1:**
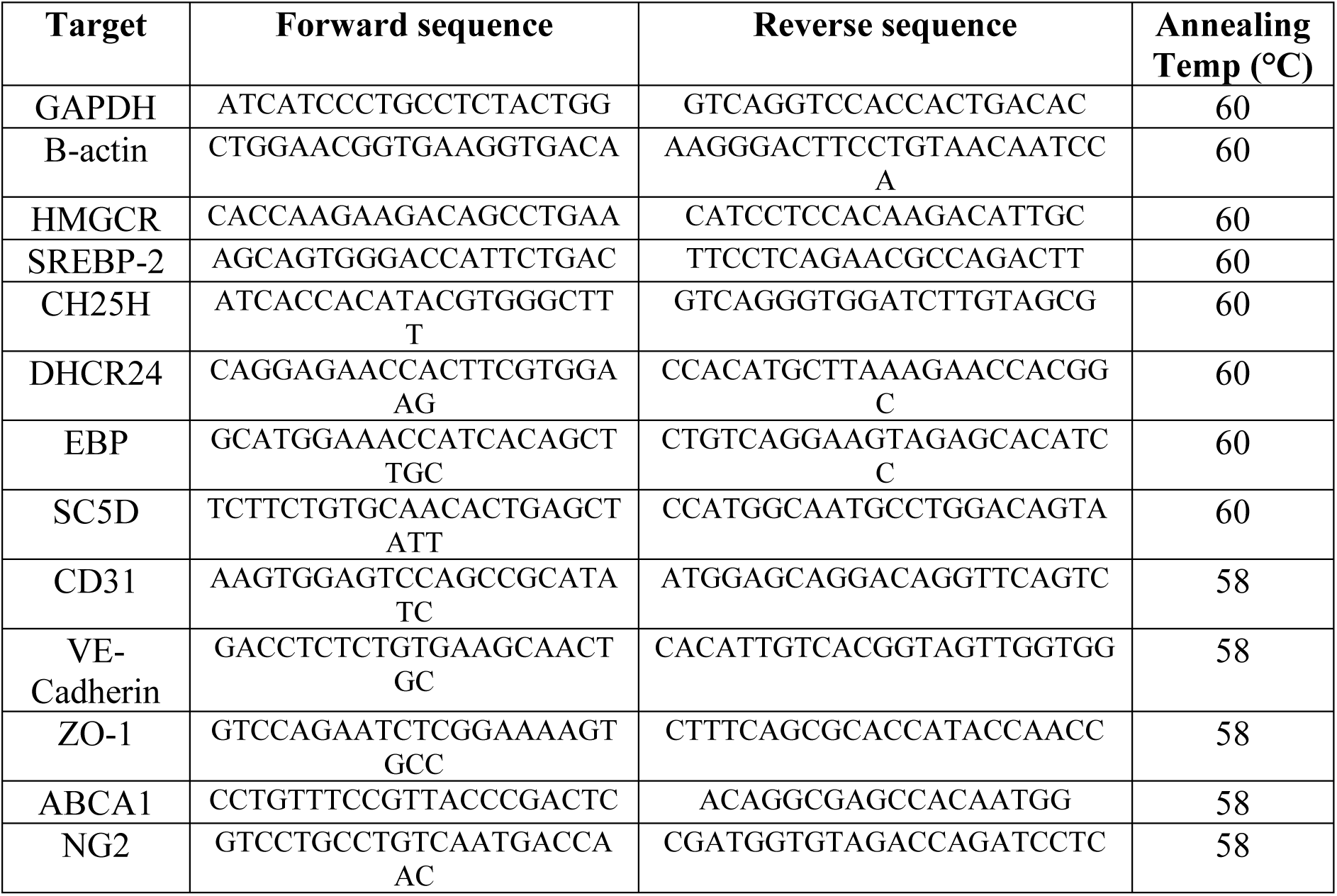
Primer sequences for qPCR analysis.

### Formalin fixed paraffin embedding tissue processing

At 40 days, vascularised organoids were fixed with 4% PFA overnight, and tissue was processed into paraffin wax at 58°C in an Excelsior AS Tissue Processor (ThermoFisher Scientific) in the histology core facility at the University of Galway. Tissue was embedded using the HistoCore Arcadia H and C (Leica), and sections were cut at 5 μm thick using a rotary microtome (Leica RM2235) and mounted on Superfrost microscopy slides (Fisher).

### Haematoxylin and eosin staining

Paraffin-embedded slides were dewaxed in xylene and rehydrated in water through a sequence of ethanol concentrations. Slides were submerged in haematoxylin solution (Mayer’s, Sigma), and an eosin counterstain was used before the tissue was dehydrated back to xylene and mounted using DPX (Sigma 06522). Images taken using the Grundium Ocus 40 slide scanner and ImageScope (Leica) software.

### Perl’s Prussian blue staining

Paraffin-embedded slides were dewaxed in xylene and rehydrated in water through a sequence of ethanol concentrations. Equal parts 10% potassium hexacyanoferrate(iii) (Merck, P8131) and 20% hydrochloric acid (Merck, 258148) were mixed just before staining. Slides were stained for 20 minutes and washed in dH2O. Slides were counterstained with nuclear fast red (Merck N3020) for 5 minutes and washed in dH2O. Slides were dehydrated through increasing ethanol concentrations and cleared in xylene for 3 minutes. Slides were mounted using DPX (Sigma 06522) and images taken using the Grundium Ocus 40 slide scanner.

### Immunohistochemistry

Cells were fixed with 4% paraformaldehyde (Thermo Fisher 043368.9L) for 1 hour at RT and washed in PBS prior to staining.

Whole organoids were fixed with 4% paraformaldehyde (Thermo Fisher 043368.9L) overnight at 4°C and washed in PBS prior to staining. Staining was carried out at 37°C.

Paraffin-embedded slides were dewaxed in xylene and rehydrated in water through a sequence of ethanol concentrations. Slides were incubated at 95°C for 15 minutes in Tris-EDTA buffer, pH 9.0, for antigen retrieval.

All samples were treated as follows. Cold blocking buffer (1% BSA, 0.3% Triton-X in PBS) was added for 1 hour. Primary antibodies were added in blocking buffer and incubated overnight at 4°C. Rabbit anti-VE-Cadherin (Cell Signalling 2500S, 1:400), rabbit anti-ZO-1 (ProteinTech 21773-1-AP, 1:500), mouse anti-CD31 (BioRad MCA1738, 1:200), mouse anti-βIIItubulin (Invitrogen PA585609, 1:500), rabbit anti-NG2 (Invitrogen PA583987 1:200). The next day, slides were incubated with secondary antibody (goat anti-rabbit AF647, Invitrogen, 1:500 and donkey anti-mouse AF488, Invitrogen, 1:500) for 2 hours at RT and mounted using Prolong Gold antifade mountant with DAPI (Thermo Fisher). Whole organoids were stained for 10 min using 1 μM Hoechst. Slides and whole organoids were imaged using the EVOS M5000 digital inverted microscope or an Olympus FV3000 laser scanning confocal microscope in the University of Galway core imaging facility. Images were processed and analysed using ImageJ and Angiotool plugin (Supplementary Table 1).

Whole organoids that were repeat stained were treated with antibody stripping solution (0.2M NaOH, 0.1% SDS) for 20 minutes at RT and then washed 3X in PBS before staining again.

### Hyperion IMC

One organoid from each treatment group was sectioned to generate 4 slides and prepared for IMC (*n*=1, 4 technical replicates). Tissue was sectioned (5 µm thick) and stained with lanthanide-conjugated antibodies as recommended by the manufacturer (Standard BioTools -https://www.standardbio.com/products/instruments/hyperion). Sections were deparaffinised, followed by antigen retrieval at 96 °C for 30 min in Tris-EDTA at pH 8.5. Non-specific binding was blocked with 3% bovine serum albumin for 45 min, followed by incubation with lanthanide-conjugated primary antibodies overnight at 4 °C, diluted in PBS with 0.5% BSA (Supplementary Table 2). Antibodies were conjugated with metals using Maxpar Antibody Labeling Kits (Standard BioTools) and were validated with positive control tissue. Slides were then washed with PBS and 0.1% Triton-X100 in PBS, and nuclear staining with iridium (1:400, Intercalator-Ir, Standard Bio Tools) was performed for 30 minutes at RT, followed by a brief (10 s) wash with ultrapure water and air-drying. Images were acquired of metal-stained tissue sections on a Hyperion imaging mass cytometer as per manufacturer’s instructions (Standard BioTools). In brief, the tissue was laser-ablated in a rastered pattern in a series of 1 µm^2^ pixels. The resulting plume of ablated tissue was then passed through a plasma source, ionising it completely into its constituent atoms. Time-of-flight mass spectrometry then discriminated the signal for each of the metal-conjugated antibodies, and images for each antibody were reconstructed based off the metal abundancy at each pixel. Staining was reviewed using MCD Viewer (Standard BioTools). Shot noise and hot pixels were removed using the IMC-Denoise algorithm. ^17^ Denoised DNA images (iridium stain) were first preprocessed using the Cellpose-3 deblur and upscale models. ^18^ Nuclear segmentation was then performed using the CellPose-SAM model, ^19^ allowing the creation of cell segmentation masks that identified cell boundaries. These cell segmentation masks were then applied to each antibody channel, generating single-cell expression data for each channel, along with the spatial context of each cell’s location in the tissue. Images were merged to present markers of interest within biological groups, and intensity expression was determined using ImageJ. Intensities were calculated as % of DNA marker expression and presented as heatmaps.

### TUNEL assay

Paraffin-embedded slides were dewaxed in xylene and rehydrated in water through a sequence of ethanol concentrations. Slides were stained using the Click-IT Plus TUNEL Assay kit AF488 (Invitrogen) as per the manufacturer’s instructions. Slides were counter stained for 10 mins using 1 μM Hoechst and mounted using Fluoromount (Merck). Slides were imaged using the EVOS M5000 digital inverted microscope and images were processed and analysed using ImageJ and Angiotool plugin.

### Statistical analysis

Data were analysed for statistical significance using GraphPad Prism (v8) and displayed as mean ±SD unless stated otherwise. Comparative analyses tests are stated for each experiment in the figure legends.

### Data availability

All raw data can be requested from the authors.

## Results

### ATV treatment causes the decrease of VE-Cadherin surface expression but increases ZO-1

Atorvastatin has been shown to disrupt endothelial cell function in vivo, thereby modelling vessel rupture in ICH pathology. To determine the effect on endothelial cells in culture, we investigated the expression of functional proteins in response to treatment.^20^ Cells were cultured in a stationary and on rotating shakers to simulate the effects of flow on endothelial cell function. ATV had no impact on cell morphology or survival in both stationary and rotating cultures, compared with DMSO controls (Figure 1A, 1B). Cells from rotating cultures exhibited gene expression changes with ATV treatment, which indicated dysregulated tight junction protein expression (Figure 1C). VE-Cadherin and NG2, an early neuro-glial marker, were overexpressed; however, this varied between cell passages sampled. Some samples showed an increase in ZO-1 expression, another tight junction protein, this was also variable and non-significant.

**Figure 1:**
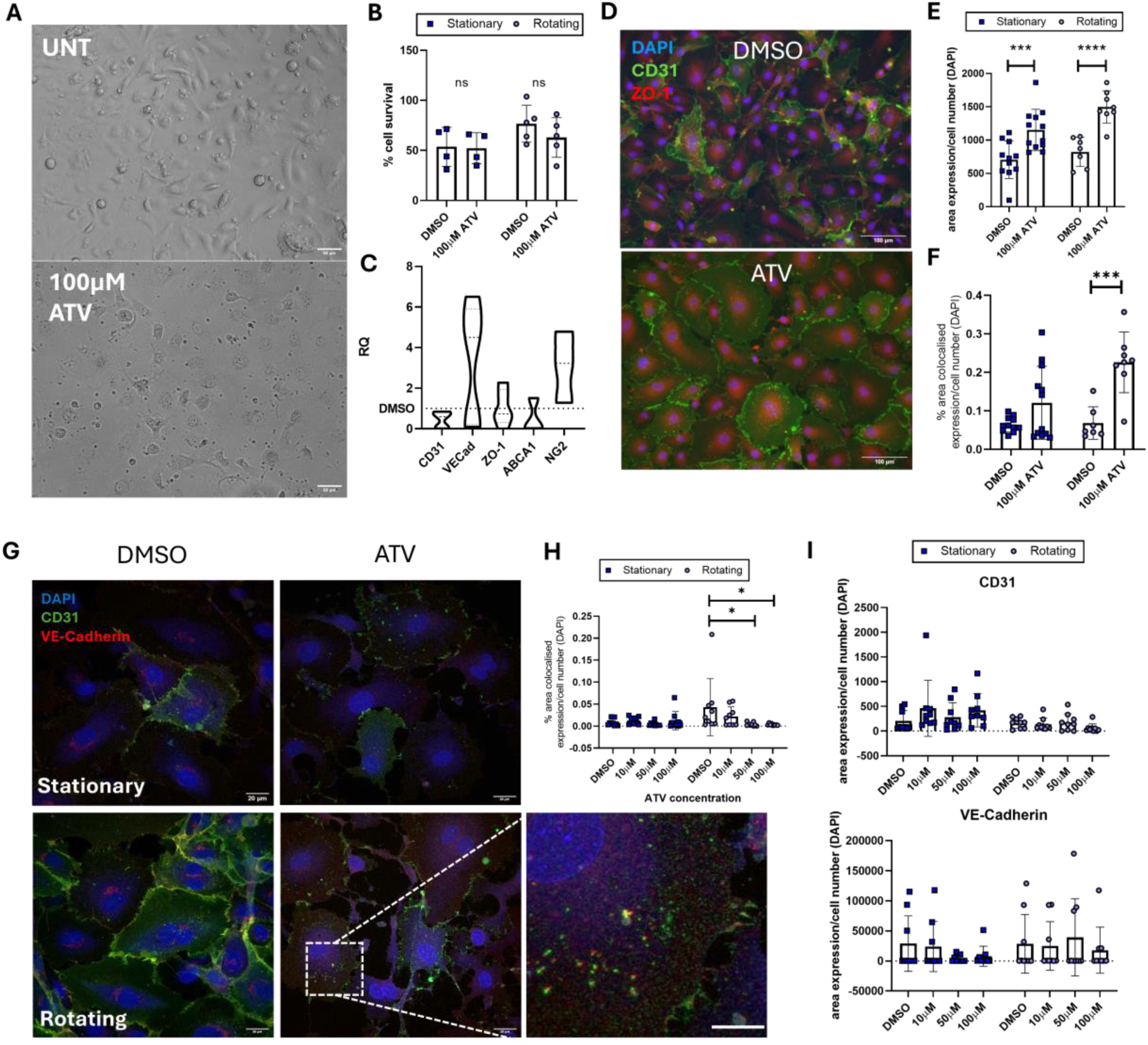
Atorvastatin treatment reduces VE-Cadherin expression on the surface of endothelial cells. (**A**) Phase contrast images of HBMECs after 24 hours of treatment with 100 μM atorvastatin (ATV). Scale bars 50 μm. (**B**) Cell survival after ATV treatment for 24 hours in both stationary and rotating culture conditions (*n*=4 independent repeats). Scale bars 100 μm. (**C**) qPCR analysis of marker expression after ATV treatment in rotating cells shows an increase in VE-Cadherin and NG2 gene expression (non-significant, One-Way ANOVA). (**D**) Fluorescent images of CD31 (green) and ZO-1 (red) cellular expression after 24 hours with ATV and DMSO control. (**E**) Total expression of ZO-1 after treatment (*n*=4 independent repeats, two-way ANOVA \*\*\**P*=0.0009, \*\*\*\**P*<0.0001) (**F**) Analysis of ZO-1 co-localised with CD31 on the cell surface when treated with ATV (*n*=4 independent repeats, two-way ANOVA, \*\*\**P*=0.0002). (**G**) Fluorescent images of CD31 (green) and VE-Cadherin (red) cellular expression after 24 hours of ATV treatment in both stationary (top panels) and rotating culture (bottom panels). Zoom inset shows the internalisation of both CD31 and VE-Cadherin from the cell surface when treated with ATV. Scale bars 20 μm. (**H**) Analysis of VE-Cadherin co-localised with CD31 on the cell surface when treated with ATV (*n*=3 independent repeats, two-way ANOVA, \**P*<0.04). (**I**) Total area expression of CD31 and VE-Cadherin as a percentage area of DAPI in both stationary and rotating cells, (*n*=3 independent repeats, no significance from analysis with a two-way ANOVA).

To determine whether this dysregulated expression was affecting protein presentation on the cell surface, ZO-1 was stained in both stationary and rotating cells, and expression intensity was significantly increased with ATV treatment (Figure 1D, 1E and 1F). VE-Cadherin was stained in both stationary and rotating cells (Figure 1G). Co-localisation of VE-Cadherin at the cell surface with CD31, an endothelial cell marker, was significantly reduced by tenfold at higher ATV concentrations (Figure 1H). However, there was no difference in the expression between culture conditions, or with the addition of ATV (Figure 1I). Data was collected from four independent repeats, and so where gene expression is not conclusively increased (Figure 1C), there may be a retention of protein at the cell surface to compensate for the loss of VE-

Cadherin. ZO-1 regulates the tension of adherens and tight junction proteins acting as an anchor,^21^ and so an increase may be due to the loss of VE-Cadherin from the cell surface. Previous studies suggest that ATV inhibits phosphorylation of these tight junction proteins and therefore, renders them incapable of localising to the membrane.^13^ We validated the findings using β-cyclodextrin, a hydrophobic inclusion complex that absorbs cholesterol, and observed no difference in the VE-Cadherin expression (Supplementary Fig. 1).

### Overall cholesterol biosynthesis is increased when treated with ATV

To determine whether ATV had an effect on the total cholesterol in the cells, filipin staining was used, and a significant increase was observed in ATV-treated cells (Figure 2A, 2B). Analysis of rotated samples showed that the transcription of the cholesterol biosynthesis gene HMGCR was significantly increased compared to DMSO controls, ultimately mitigating inhibition by ATV (Figure 2C). Other biosynthesis genes (SREBP-2 and SC5D) were also increased however this was not found to be significant from DMSO treated controls. CH25H and DHCR24 were not detected in any samples.

**Figure 2:**
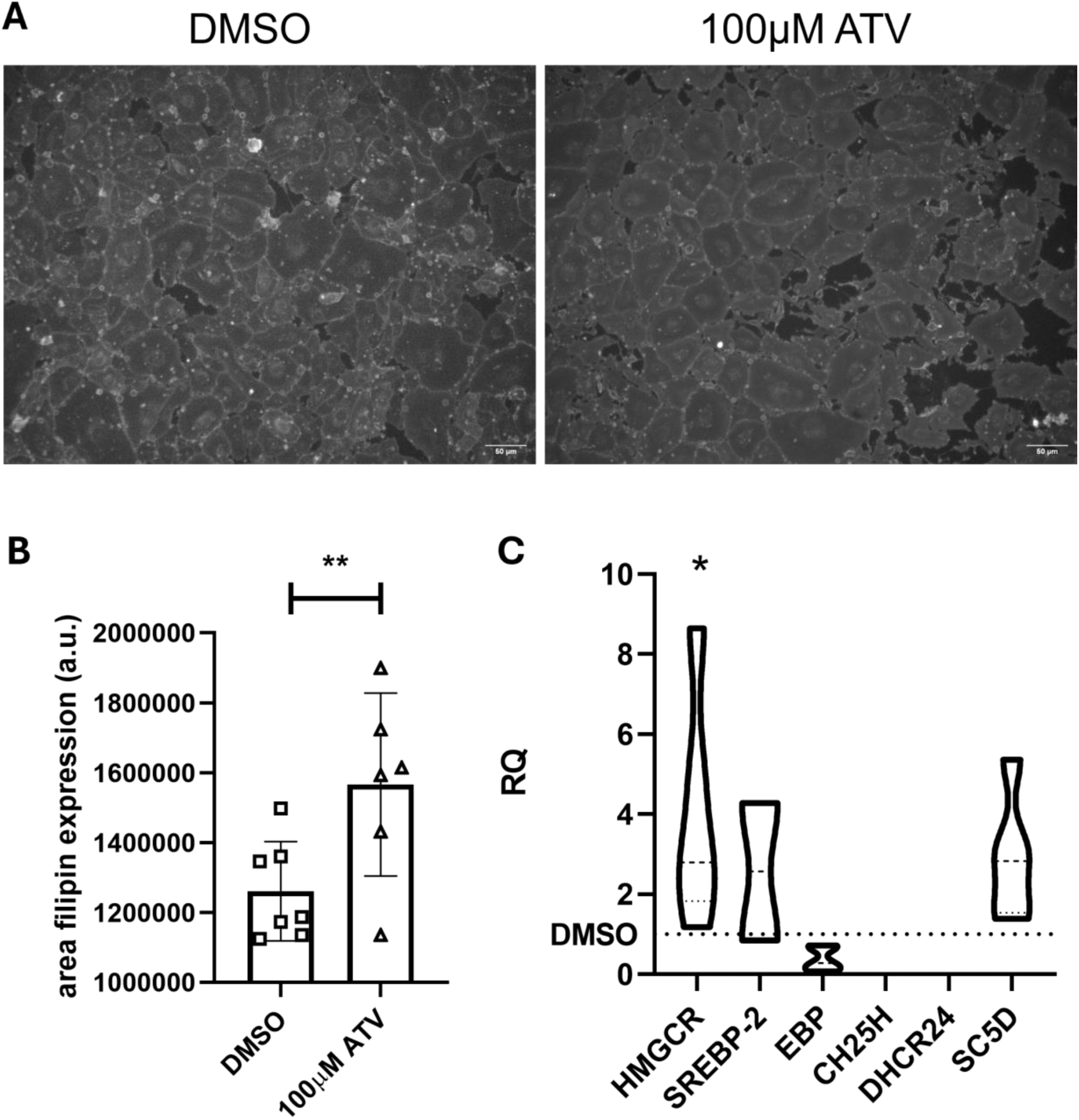
Cholesterol levels increase HBMECs after ATV treatment. (**A**) Representative images of filipin-stained HBMECs, after 24hours of drug treatment. Scale bars 50 μm. (**B**) Filipin expression as a percentage of area from 6 ROIs in 2 wells across 2 independent repeats after 24 hours with ATV, ** *P*=0.0081, *t*-test. (**C**) qPCR analysis of rotated HBMECs after 24 hours of drug treatment shows a significant increase in HMGCR expression (*n*=5 independent repeats, * *P*=0.0323 One-Way ANOVA).

### ATV disrupts the tube formation property of endothelial cells irreversibly

To determine whether the loss of tight junction protein VE-Cadherin had a detrimental effect on the functional properties of endothelial cells, we conducted a tube formation assay. Endothelial cell tubes that were treated with ATV demonstrated a significant disruption in tube morphology (Figure 3A), and a significant reduction in the length of tubes (Figure 3B). Cells that were treated with ATV before tubes formed were unable to form tubes after 24 hours, implying a short-term irreversible effect (Figure 3B). No cytotoxic effect was observed (Figure 3C).

**Figure 3:**
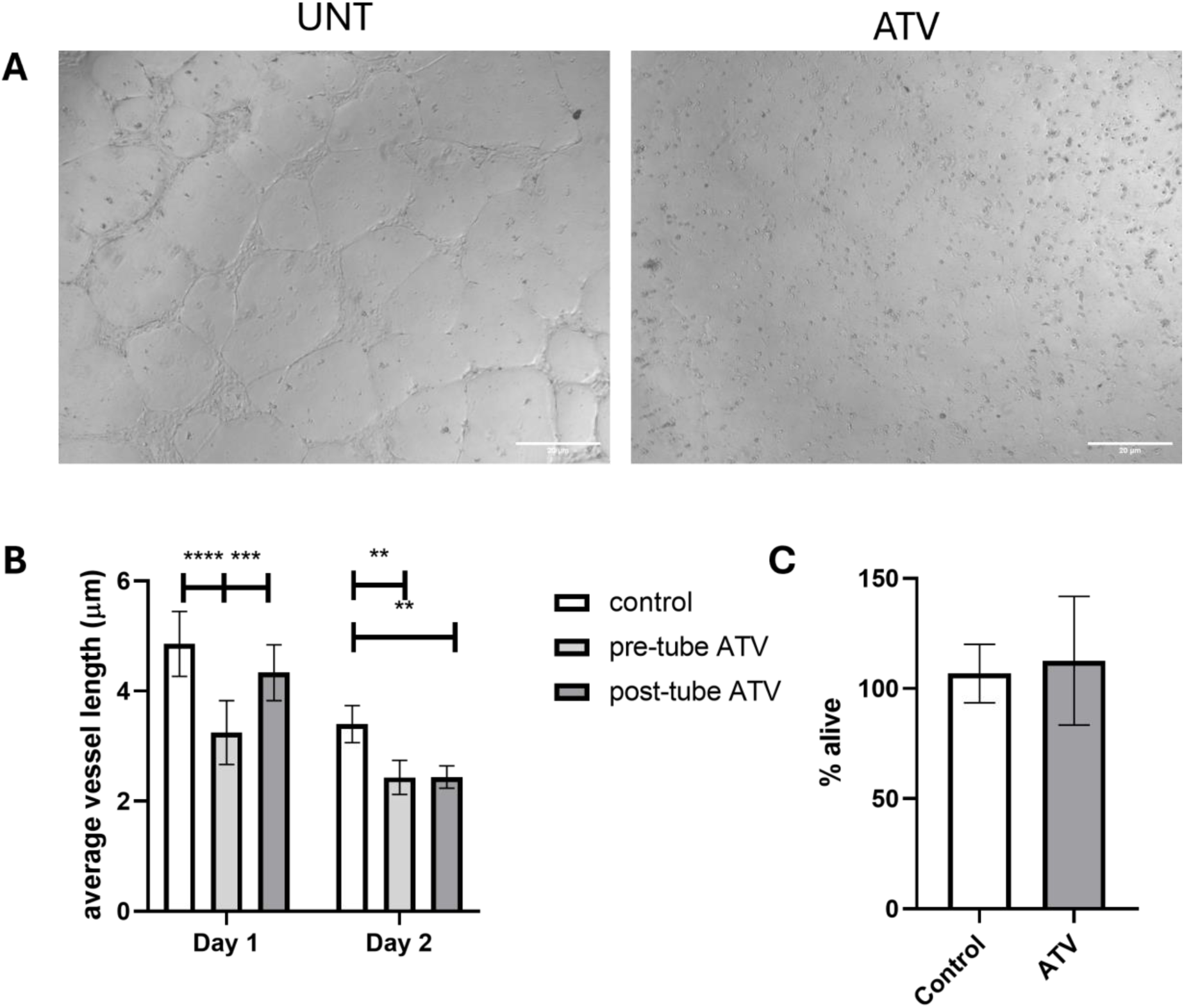
Atorvastatin disrupts endothelial tube formation. (**A**) Phase images of tube formation in untreated and ATV-treated HBMECs. Scale bars 20 μm. (**B**) Analysis of the network parameter average vessel length shows a significant reduction with ATV treatment, both pre-tube formation and post (One-way ANOVA with Tukey’s *post hoc* multiple comparisons test ** *P*=0.003, *** *P*=0.0007, **** *P*<0.0001). (**C**) Cytotoxicity analysis of ATV treated tubes showed no difference from controls (*n*=3, *t*-test).

### In 3D organoid tissues, ATV causes the disruption of endothelial cell networks and a decrease in VE-Cadherin expression

Of interest for the application of ATV to ICH modelling in organoids, we investigated whether ATV could disrupt the endothelial cell networks that form on vascularised cerebral organoids (V-COs).^9^ V-COs were treated with ATV for 24 hours after 40 days of culture, and staining revealed that the network coverage of CD31-positive cells and associated VE-Cadherin expression was disrupted on the surface of the organoid (Figure 4A). When cell marker expression was quantified as a percentage of DAPI, there was a significant reduction in VE-Cadherin expression by more than 50% in ATV-treated V-COs compared to DMSO controls (Figure 4B). This was validated in organoids from four separate batches (Figure 4C) and in organoids derived from an additional cell line (Supplementary Fig. 2). However, this loss of expression did not translate into significant changes in the CD31 network morphology, as determined by Angiotool analysis of the number of vessel end points (fragmentation) and the average vessel length in μm (Figure 4D) but did result in significantly shorter vessels detected by VE-Cadherin staining (Figure 4E). Gene expression analysis through qPCR revealed that ATV-treated V-COs exhibited no change in expression of cell markers such as ZO-1 and functional ABCA1, and an overall decreased expression of CD31, VE-Cadherin and NG2, although not significant (Figure 4F), opposite to what was observed in 2D cultures (Figure 1C). An upregulation of the downstream cholesterol biosynthesis proteins EBP and SC5D and a decrease in HMGCR, not previously observed in 2D cultures, suggest that the neural tissue and 3D behaviour of endothelial cells drives the increase in expression (Figure 4G).

**Figure 4:**
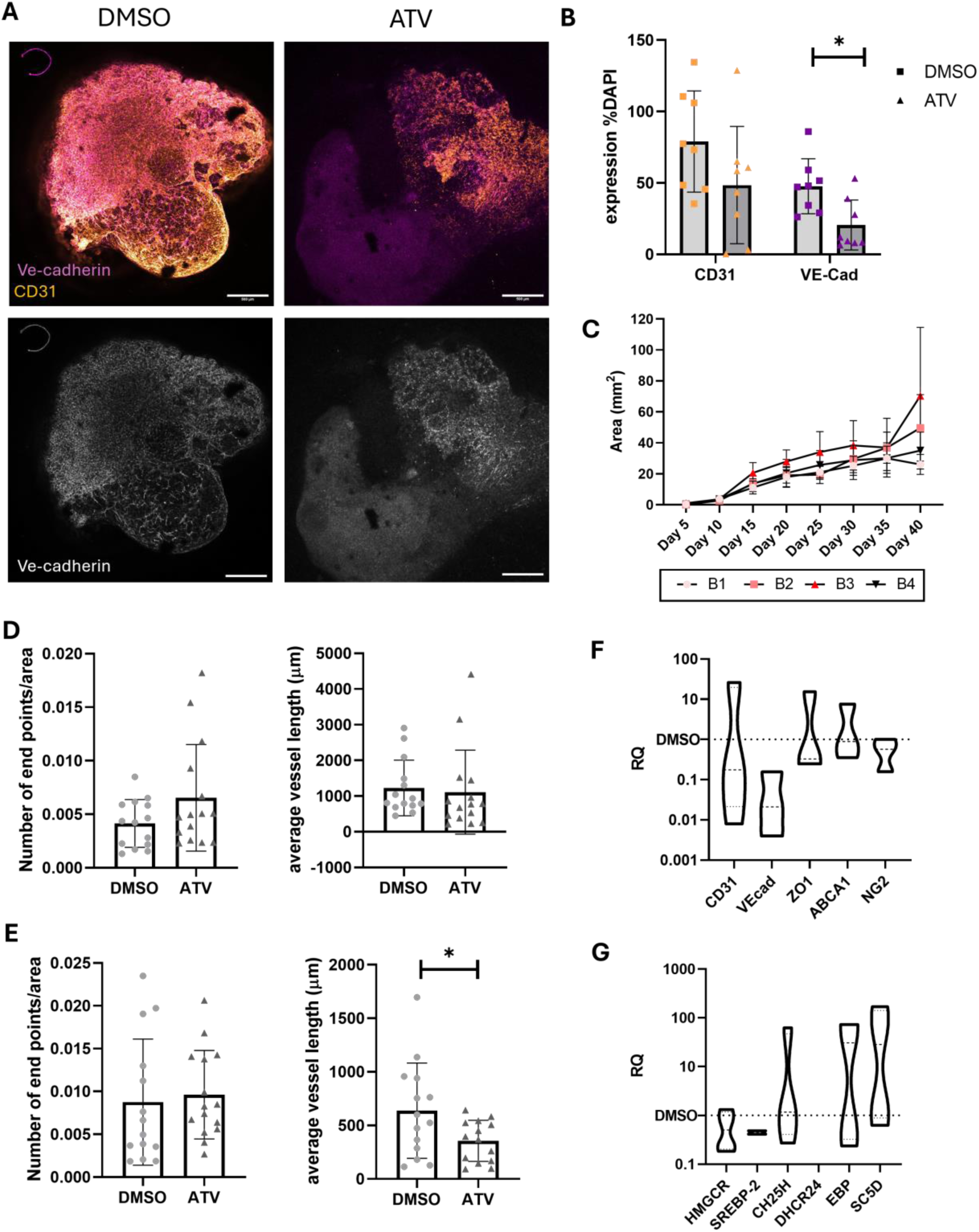
**ATV treatment reduces VE-Cadherin expression on the surface of organoids.**(**A**) Representative confocal images of whole organoids treated with ATV for 24 hours and the loss of VE-Cadherin expression from the surface. Scale bars 500 μm. (**B**) VE-Cadherin, not CD31 (*P*=0.091), expressed as a percentage of DAPI was significantly reduced with ATV treatment compared to DMSO controls (two-tailed *t*-test, * *P*=0.011). (**C**) Size progression for 4 batches of organoids that were treated with ATV at day 40. (**D**) Vascular metrics were unchanged for CD31 (*n*=15 organoids from 4 batches, non-significant *t-*test) with slightly more endpoints, signifying single cells. (**E**) Angiotool analysis of VE-Cadherin staining revealed shorter overall vessel length (*n*=15 organoids from 4 batches, *t-*test, \**P*=0.0389)). (**F**) qPCR analysis of ATV-treated organoids showed a reduction in VE-Cadherin RNA expression; however, other markers of endothelial function are unchanged (One-Way ANOVA, *n*=5 organoids from 2 batches). (**G**) ATV treatment increased the expression of some cholesterol biosynthesis markers compared to DMSO controls, opposite to what was observed in HBMECs in 2D (One-Way ANOVA, *n*=5 organoids from 2 batches).

### ATV treatment affects internal extracellular matrix structure

Because ATV treatment disrupts the expression of VE-Cadherin on the organoid surface, we sought to determine what occurred within the tissue. Organoids were sectioned and stained, and haematoxylin and eosin histology showed the extent of the heterogenous tissue patterning within a single organoid (Figure 5A). Both DMSO and ATV-treated organoids show regions of neural rosette formation (green arrow), lumenised ventricular spaces (blue arrow) and less cell-dense, more vascularised regions (red arrow). Endothelial cells (CD31) and VE-Cadherin were stained for in internal slices and no differences in expression were detected (Figure 5B). Likewise, βIIItubulin was also unaffected by ATV treatment, assuming a limited effect on the neural tissue (Figure 5C). These data may indicate that ATV has a local effect on VE-Cadherin on the surface of endothelial cells, and, because there is no perfusion of the tissue, there is insufficient internal exposure to ATV. To confirm the breadth of impact of ATV and blood treatment on internal tissue, organoids were analysed using Hyperion Imaging Mass Cytometry (IMC). Four slices from one organoid of each group were imaged (Figure 5D) and intensity of each channel was expressed as a % of DAPI (Figure 5E). Observations from images revealed the presence of microglial marker ionised calcium-binding adaptor molecule (IBA1), previously unobserved by immunohistochemistry. The extracellular matrix (ECM) was largely formed of collagenIV and fibronectin, not found in Geltrex and both important for brain development. Fibronectin is upregulated by activated astrocytes (glial fibrillary acidic protein - GFAP) and is associated with vascular cells (aquaporin 4 -AQP4) and with increased phosphorylated NFκB (pNFκB). Expression of fibronectin, collagenIV and smooth muscle actin (SMA) was decreased in ATV organoids with blood. There was no effect of ATV on neural markers, or proliferating cell markers (nestin, NeuN, vimentin, SOX2, SOX10, Ki67 or phosphorylated ERK). Cells were segmented with a 1 μm radius (Supplementary Fig. 3) and expression of cell-associated markers was normalised to the 99th percentile (Figure 5F). The summary heatmap of the cell-segmented data emphasises the trends observed in Figure 5E, notably an increase in cleaved caspase 3 expression with ATV treatment and a potential decrease in CD31 expression with blood treatment.

**Figure 5:**
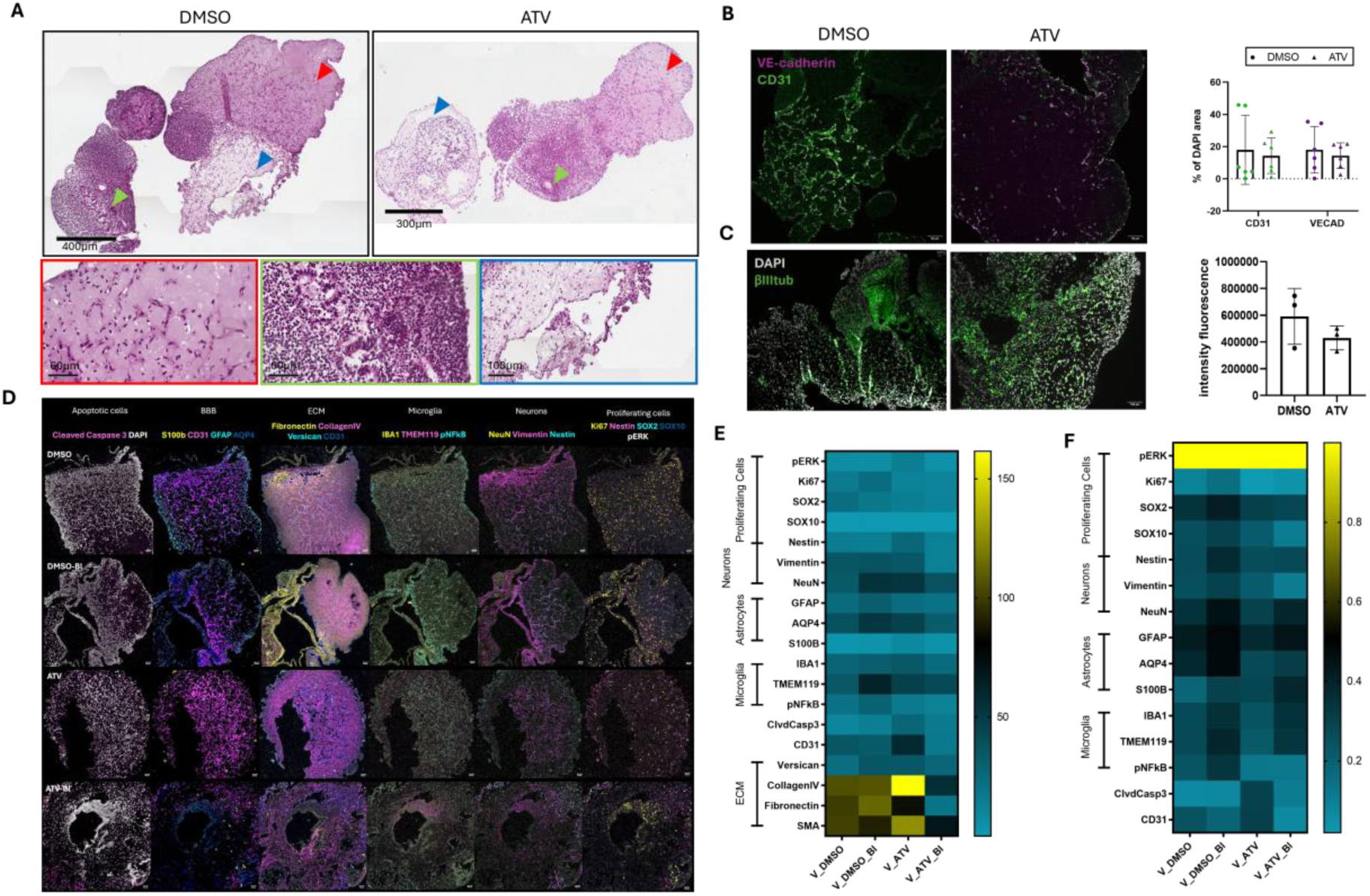
Internally, ATV has no effect on tight junction expression but in combination with blood, depletes the extracellular matrix. (**A**) Haematoxylin and eosin staining of internal slices from organoids treated with DMSO or ATV. Different tissue morphology is visible throughout both tissues. Scale bars 300 μm. (**B**) Immunostaining for CD31 (green) and VE-Cadherin (magenta) show no difference in intensity of expression with ATV treatment. Scale bars 50 μm. (One-Way ANOVA, *n*=6 organoids, three independent batches). (**C**) No effect significant effect on βIIItubulin expression with ATV treatment, although there may be a decrease. Scale bars 100 μm. (*t*-test, *n*=3 organoids). (**D**) Representative images from Hyperion IMC analysis of 4 slices of each organoid from each group. Scale bars 50 μm. BBB: blood-brain barrier, ECM: extracellular matrix, GFAP: glial fibrillary acidic protein, AQP4: aquaporin 4, IBA1: ionised calcium-binding adaptor molecule 1, pNFκB: phosphorylated nuclear factor kappa-light-chain-enhancer of activated B cells, NeuN: neuronal nuclei, Ki67: antigen Kiel 67, pERK: phosphorylated extracellular signal-related kinase. (**E**) Heatmap of intensity expression as a % of DAPI from each channel after denoising showing the combined effect of ATV and blood at the expression of extracellular matrix proteins. (**F**) Heatmap of intensity expression normalised to 99^th^ percentile within segmented cells.

### Exposure to human blood does not exacerbate the loss of vascular networks on the surface of the organoid

Whole human blood was added to induce tissue damage similar to that seen in clinical ICH. Whole blood contains a host of factors that are neurotoxic and stimulate an innate immune response.^22^ When blood was added to the organoid media after 40 days of culture, it was visible on the surface of the organoid (Figure 6A). More blood was visibly adhered to the surface of organoids that had been treated with DMSO compared to ATV, this was only observational, as when tissue was fixed, blood was subsequently washed off. Whole organoid staining for CD31 and VE-Cadherin (Figure 6B) shows that blood-treated organoids had no additional effect on the reduction of VE-Cadherin-positive vessel length (Figure 6C). Vascular metrics of CD31 positive staining was unchanged by blood addition to culture (Figure 6D). Interestingly, blood addition to ATV-treated organoids showed an increase in VE-Cadherin expression back to control levels (Figure 6E, non-significant difference between groups). This could be due to blood factors that increase endothelial cell function.

**Figure 6:**
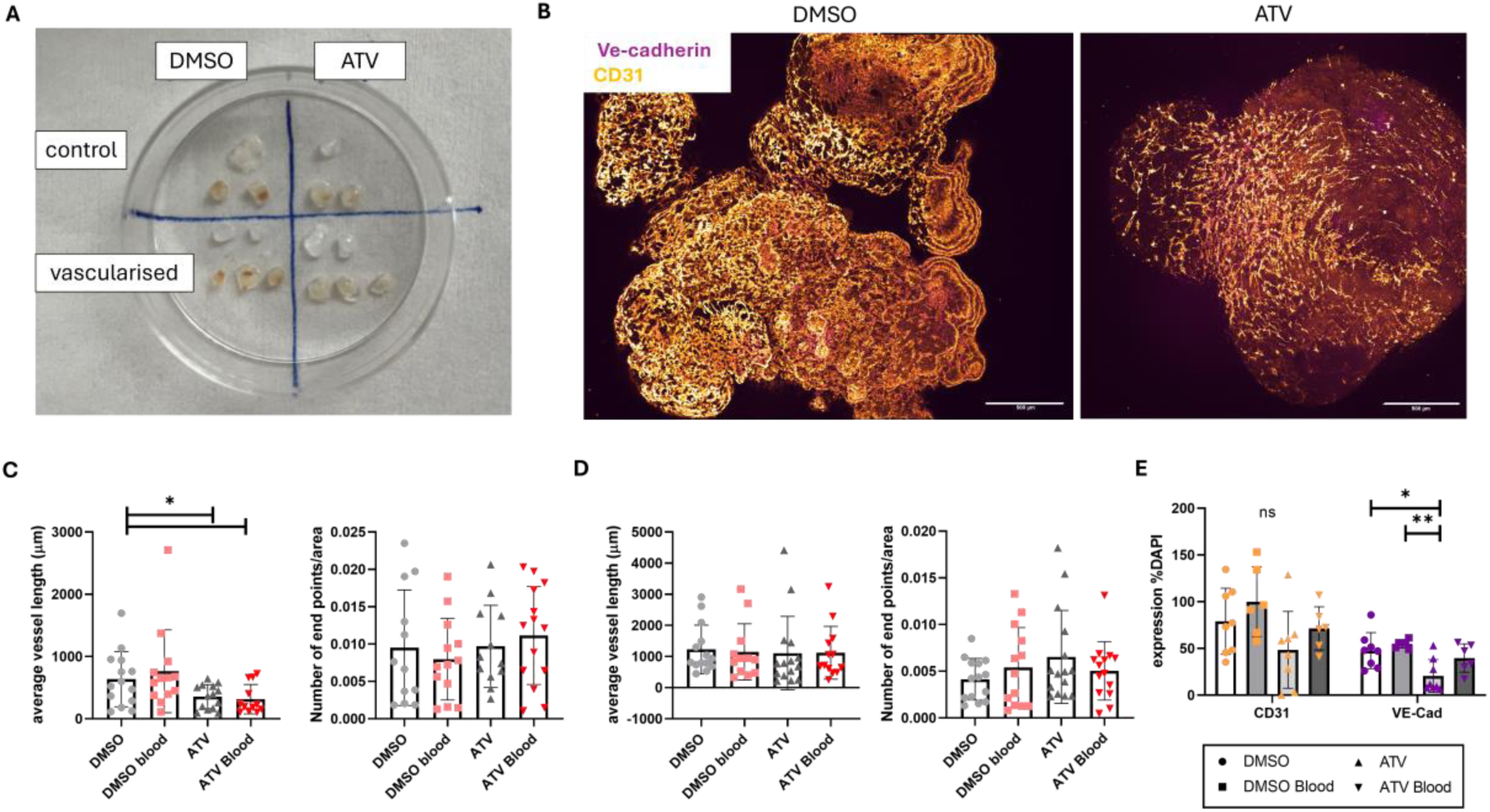
Human blood exposure does not exacerbate vessel loss on the organoid surface. (**A**) Representative image of 40-day organoids that have been treated with whole human blood for 4 hours, prior to washing and fixing steps. (**B**) Confocal images of the surface CD31 networks and VE-Cadherin expression in organoids post DMSO and ATV treatment, incubated in whole human blood, show no morphological differences. Scale bars 500μm. (**C**) Angiotool analysis of VE-Cadherin networks (unpaired *t*-tests \**P*=0.03) (**D**) CD31 networks shows no significant differences when organoids are treated with blood (*n*=12 organoids from 4 batches, One-way ANOVA). (**E**) Expression intensity analysis shows no significant differences in CD31 expression with ATV and/or blood; however, despite the decrease in VE-Cadherin expression with ATV treatment, this increased in organoids treated with blood (One-Way ANOVA with Tukey’s post hoc multiple comparison analysis; * *P*=0.0109, ** *P*=0.0026).

### Human blood exposure causes overall tissue damage and neuroinflammation

Perl’s Prussian blue iron stain detected iron (blue staining, black arrows) on internal slices of the organoid tissue (Figure 7A) and perhaps some insoluble hemosiderin deposits (brown staining, red arrow) from the lysed red blood cells. This suggests that the blood permeates the ‘leaky’ tissue and reaches some internal cells. The location of iron staining did not correspond with any local morphological differences observed with H&E staining (top right panel). However, some iron and haemosiderin staining appears in a similar location with TUNEL-positive apoptotic cells (bottom right panel, white arrows, *n*=11/18).

**Figure 7:**
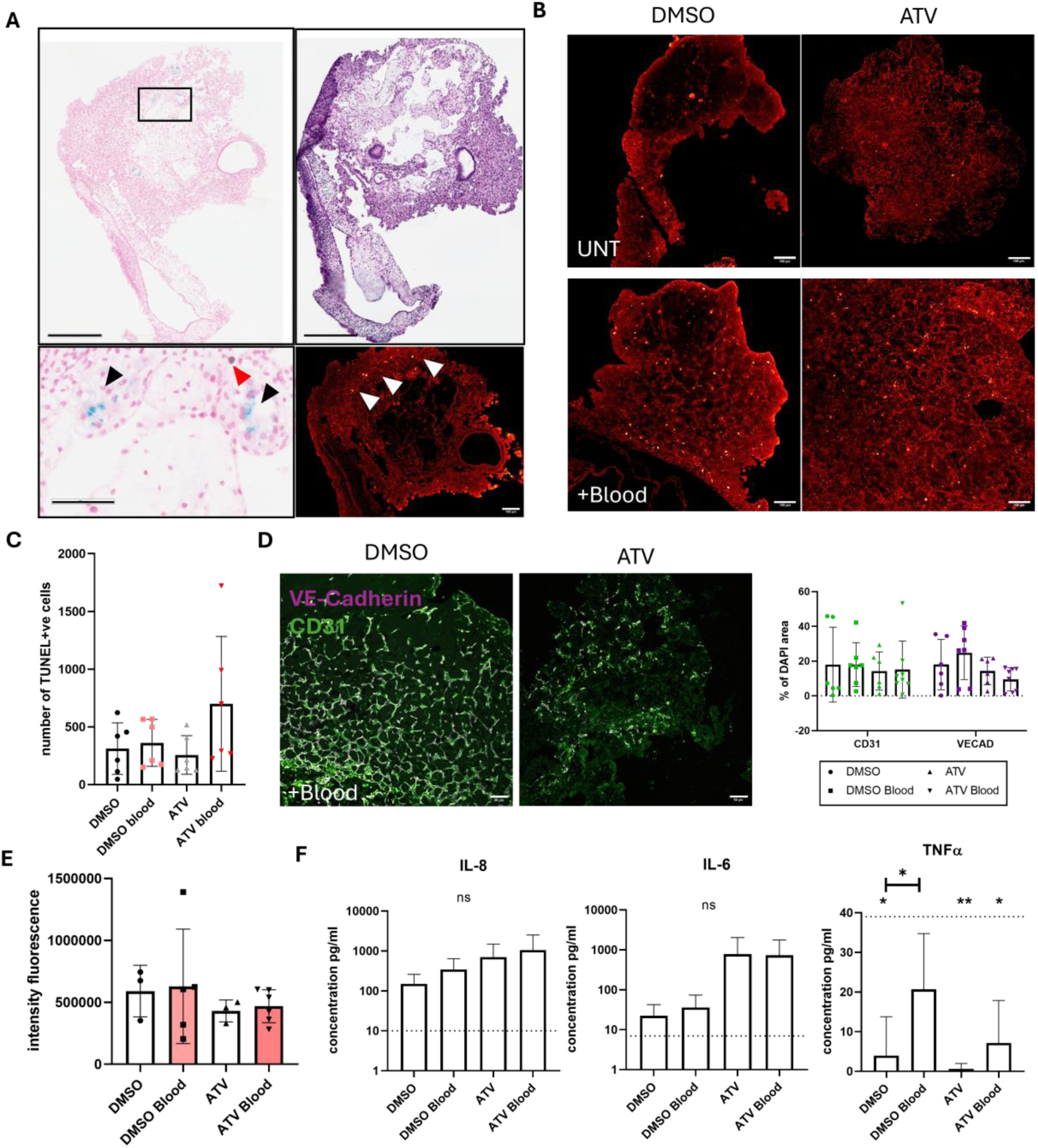
Blood penetrates the tissue, causing cellular damage and initiating an immune response. (**A**) Perl’s Prussian blue staining for iron identifies iron deposits within the organoid tissue (black arrows, scale bar 60μm); however, no morphological differences were observed with H&E staining (right panel, scale bar 300μm). Areas positive for iron staining were often associated with TUNEL-positive apoptotic cells (white arrows, *n*=11/18, scale bar 100μm). (**B**) Representative images of slices from organoids treated with DMSO and ATV with and without blood and stained for TUNEL-positive apoptotic cells (*n*=3 from 2 independent batches, scale bar 100μm). (**C**) Quantification of the TUNEL staining shows an increase in the ATV+Blood treated organoids compared to ATV alone, or DMSO+Blood, this was non-significant (*n*=6, One-Way ANOVA). (**D**) Blood treated organoids showed no difference in internal networks of CD31 and VE-Cadherin compared to untreated groups. Scale bars 50μm. (**E**) Blood treatment showed no significant effect on βIIItubulin expression. (**F**) ELISA analysis of the inflammatory markers released into the media after 4 hours incubation with whole blood. Dotted line represents whole blood control (*n*=3 independent batches, One-Way ANOVA with Tukey’s post hoc analysis \**P*=0.0258, One-Way ANOVA with Dunnett’s multiple comparisons to whole blood \**P*<0.011, \*\**P*=0.0053).

TUNEL analysis of apoptosis revealed that ATV in the V-COs had no effect on cell death (Figure 7B), the toxicity of blood in some slices was doubled in ATV-treated V-COs (Figure 7C). This, taken with data from 8A, perhaps implies that the ATV-treated V-COs were ‘leakier’ than DMSO controls due to the loss of integrity in the surrounding vascular network for protecting the neural tissue. Blood treatment does not seem to affect the TUNEL-positive apoptotic cells in DMSO controls, perhaps because this analysis is from internal slices rather than local contact effects on the surface of the organoid. To confirm this, we observed that blood treatment did not affect the CD31 and VE-Cadherin networks within the tissue (Figure 7D) or the intensity expression of βIII tubulin (Figure 7E), implying that the TUNEL-positive cells are more likely to be neural than vascular.

Assessment of the neuroinflammatory markers released into the media after culture with whole blood was performed by ELISA (*n*=3 batches of organoids) (Figure 7F). IL-1β and IL-10 were undetected in all samples. Increased levels of interleukin (IL)-6 and IL-8 were detected after 4 hours in all organoid samples compared to whole blood, however, non-significant. ATV treatment appears to cause a 20X increase in IL-6 detection compared to DMSO controls.

Blood treatment increases IL-8 release compared with DMSO/ATV treatment alone. TNF-α was observed to decrease compared to levels in the whole blood control, significantly in DMSO, ATV and ATV Blood samples, and blood significantly increased TNF-α release in DMSO-treated groups.

## Discussion

Here we have shown that vascularised human brain organoids may offer a platform for investigating intracerebral haemorrhage, and blood insult in the brain. Recent vascularisation of cerebral organoids has allowed for the in vitro modelling of the human cerebrovasculature, and there are key aspects of the BBB replicated in this platform that make it suitable for the interrogation of ICH. For the first time, this provides a wholly human platform for pre-clinical investigation and opens the door for modelling in patient-derived tissues for stroke. This model platform offers a human complement to the animal models as an adaptable alternative with multi-systemic complexity. We have shown that atorvastatin may result in vascular weakness through depletion of VE-Cadherin expression at the cell surface and a resulting loss of cellular network function. We observed for the first time with IMC the presence of IBA1 positive cells, and key microglial markers, and noted the loss of fibronectin, collagenIV and versican in the presence of ATV and blood. This study also revealed the close spatial association of nestin, phosphorylated NFκB and AQP4 with the CD31 positive vasculature, and further detailed investigation will reveal the extent of the impact of vascularisation on the neural tissue differentiation, maturation and function. This translates to 3D models, and although the mechanism is not fully understood, may lead to a controllable, local effect on cells when tissue is perfused. We also show here the V-CO response to human whole blood, increased iron presence within the tissue, increased apoptotic cell death and a stimulated innate immune response from cerebral and vascular tissues.

There are limitations and optimisations that can be made to the model. The model is not perfused and accurate; directional perfusion still eludes much of the organoid community. There is a drive to install organoid microphysiological systems into microfluidic chips to control the flow of media and drug delivery; however, differing vascularisation techniques and the size of the organoid tissue require bespoke chips and optimised incorporation steps.^3^ We hope to overcome this particular limitation to exert a localised effect of ATV within the vascular networks at the depth of the tissue, more closely mimicking the vascular weakness effect of haemorrhagic stroke in future studies. There is also wide heterogeneity in tissue development between organoids and batches, and the iPSC line has a definitive effect on the final microtissue produced.^9^ The overall aim of this work is to ensure the model is as adaptable as possible, so that non-specialist labs may adopt the technique and streamline pre-clinical ICH investigation.

Unexpectedly, total cholesterol levels proved difficult to measure in a 3D tissue of multiple cell types, however if there is more cholesterol with decreased gene expression in 2D cultures and the opposite in the 3D then it may mean there is less cholesterol present in the organoid. This would correlate with what is observed clinically with ATV treatment^23^ and in vivo.^10,12^ β-cyclodextrin was not investigated in the organoid because evidence from 2D cultures suggested the opposite effect to ATV however such treatment would confirm this hypothesis by reducing cholesterol directly. Alternatively, evidence suggests that ATV can also actively inhibit tight junction function by causing the internalisation of VE-Cadherin, supported by other in vivo investigations that suggest that reversing the effect on VE-Cadherin can prevent haemorrhage-like phenotypes.^13,24^

We observed that blood treatment reversed the decrease in VE-Cadherin expression caused by ATV alone. Future study would include isolating CD31+ cells after blood treatment to investigate gene expression changes and understand how cell function and maturation is affected. However, it is also essential to understand the factors in the whole blood samples that are responsible for this response as physiologically, blood flow exerts influence over endothelial cells through mechanical force such as shear and can alter polarity, transduction, migration and signalling.^25,26^ It is unclear which significant signalling factors in whole blood influence this relationship, so further study is needed.

Blood treatment stimulated an immune response from the neural and vascular cells within the system. Without a specific immune component to the organoid model, these cytokines were limited. There are culture strategies to incorporate primary, or iPSC-derived microglia into the organoid cultures^27^ However, this has not yet been attempted with V-COs. IL-8 is produced by endothelial cells and has a role in angiogenesis. It acts as a chemotactic marker, recruiting peripheral neutrophils to the injury site. It has been observed clinically in both the blood and cerebrospinal fluid of ischemic stroke patients. IL-8 was increased in all organoid cultures compared to human whole blood controls and this may be due to the specific presence of endothelial cells. Clinically, increased peripheral IL-6 is associated with worse functional outcomes^28^ whereas higher, localised IL-6 within the haematoma contributes to a good ICH outcome (INFLAME-ICH) as it has both pro- and anti-inflammatory roles. We observed increased release with ATV exposure in the V-CO model. It may be that IL-6 is released in response to vascular disruption, as there was little effect of blood in both DMSO and ATV groups.

This model can provide a platform for investigating blood neurotoxicity, induction of cerebrovascular weaknesses, and investigating action mechanisms for preventative therapeutics against blood-induced pathology in a human system. Our current pre-clinical pipeline for intracerebral haemorrhage investigation lacks complex, tunable human models beyond functional assessment, peripheral clinical samples and post-mortem tissues. With this study, we offer a complementary system in an attempt to advance the effective clinical translation of pre-clinical research.

## Author Contributions

SC is responsible for the project conceptualization, methodology, investigation, data acquisition, analysis, and visualisation, funding and the writing of the original manuscript, and supervision for ES and NC. SS, ES, NC and EJL contributed to the investigation and data acquisition. MH and AC are responsible for the data acquisition, analysis and visualisation. JFN contributed resources. KNC, DM and ML are responsible for supervision. ML is responsible for conceptualisation, provision of resources, and project administration. All authors were involved in the reviewing and editing of the final manuscript.

## Acknowledgements

The authors would like to thank Dr Karolina Salcuite, Dr Keith Madden and Dr Jamie Reilly in the University of Galway Technology Services Directorate for facility use and technical assistance. We thank the Flow Cytometry and the Bioimaging Core Facilities at the University of Manchester for helping with the imaging mass cytometry work performed in this study. The imaging mass cytometer used within the study was purchased through a BBSRC Alert18 award (BB/S019324/1 to K.N.C.).

