## Supplementary File for "Atorvastatin-induced vascular weakness models brain haemorrhage in vascularised cerebral organoids"

*Supplementary Information*

S-iPSCs (STEMCELL TECHNOLOGIES Healthy Control Human iPSC Line SCTi003-A, Cat# 200-0511, Female)

Beta-cyclodextrin hydrate (Thermo Scientific A14529.06) was reconstituted in a 1:1 mix of distilled water and DMSO to generate a stock solution.  Treatment was carried out as with ATV, using 1:1 DMSO:water control.

*Angiotool methodology*

| | |
|---|---|
| Width | 2048 |
| Height | 2048 |
| Low Threshold | 25 |
| High Threshold | 255 |
| Vessel Thickness | 1 |
| Small Particles | 3 |
| Fill Holes | 5 |
| Max Hole Level | 1.5 |
| Min Boxness | 0.09375 |
| Min Area Length Ratio | 16 |
| Skeletonizer | Thorough |
| Max Skeleton Steps | 25 |
| Scaling Factor | 1 |
| Transformed Colors | No |

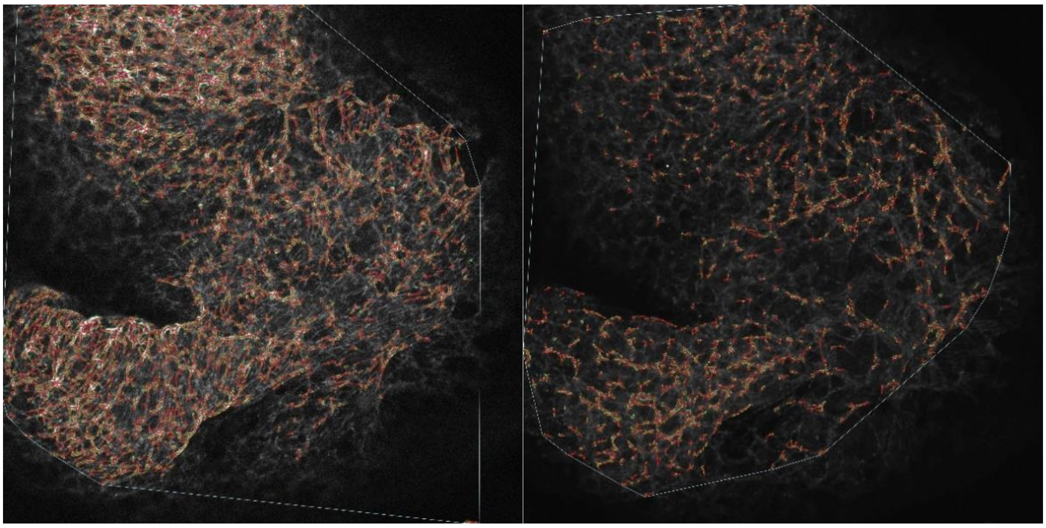

Representative images of Angiotool analysis of CD31 (left) and VE-Cadherin (right)

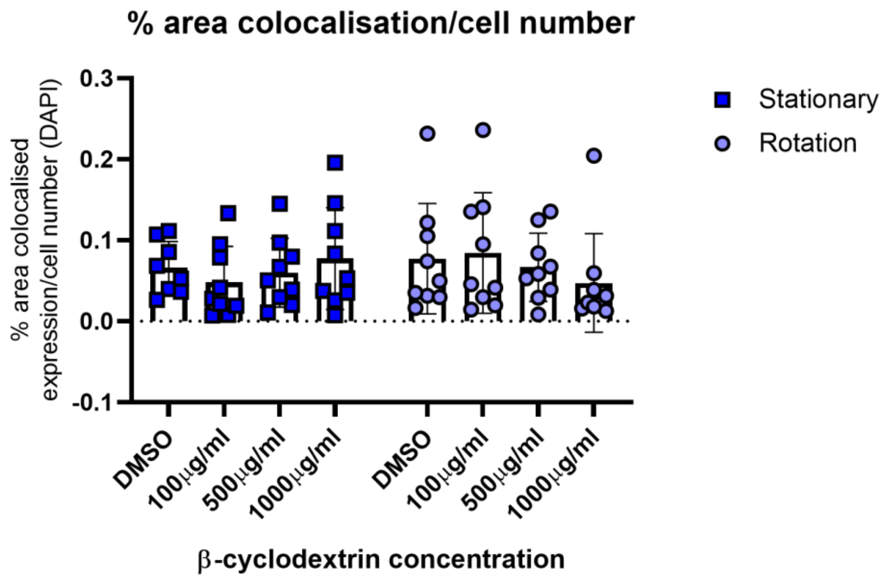


**Supplementary Figure 1:** βcyclodextrin had no effect on the co-localised expression of VE-Cadherin with CD31 in both stationary and rotating cultures (n=3 independent replicates, Two-Way ANOVA)

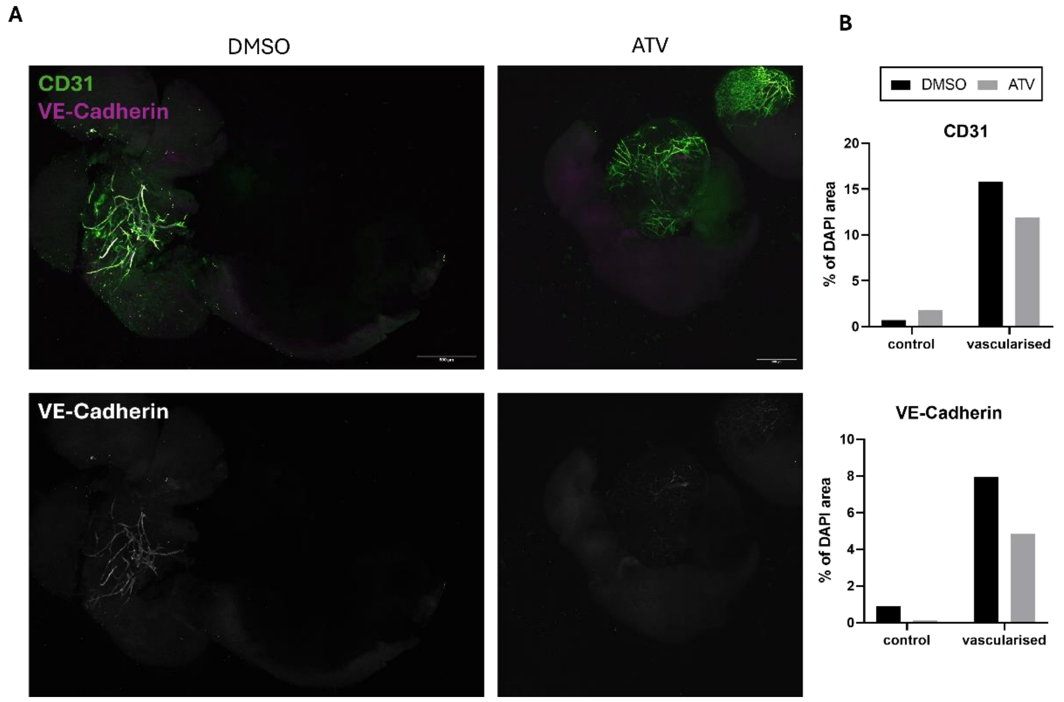


**Supplementary Figure 2: ATV treatment reduced VE-Cadherin expression on the surface of S-iPSC derived organoids.** (**A**) Representative confocal images of whole organoids treated with ATV for 24 hours and the loss of VE-Cadherin expression from the surface. Scale bars = 500µm. (**B**) Both CD31 and VE-Cadherin expression appear to decrease with ATV treatment.

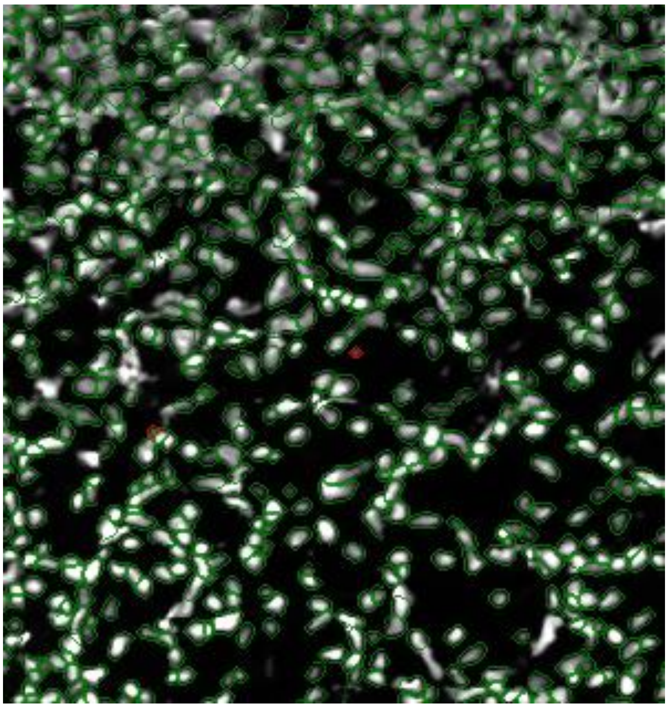

**Supplementary Figure 3: Cell segmentation for IMC images.** Mask overlay for cells (green) based on a 1 μm circumference from the DAPI positive nucleus. Objects too small to be recognised as cells are masked in red.
